# Accuracy and efficiency of germline variant calling pipelines for human genome data

**DOI:** 10.1101/2020.03.27.011767

**Authors:** Sen Zhao, Oleg Agafonov, Abdulrahman Azab, Tomasz Stokowy, Eivind Hovig

## Abstract

Advances in next-generation sequencing technology has enabled whole genome sequencing (WGS) to be widely used for identification of causal variants in a spectrum of genetic-related disorders, and provided new insight into how genetic polymorphisms affect disease phenotypes. The development of different bioinformatics pipelines has continuously improved the variant analysis of WGS data, however there is a necessity for a systematic performance comparison of these pipelines to provide guidance on the application of WGS-based scientific and clinical genomics. In this study, we evaluated the performance of three variant calling pipelines (GATK, DRAGEN™ and DeepVariant) using Genome in a Bottle Consortium, “synthetic-diploid” and simulated WGS datasets. DRAGEN™ and DeepVariant show a better accuracy in SNPs and indels calling, with no significant differences in their F1-score. DRAGEN™ platform offers accuracy, flexibility and a highly-efficient running speed, and therefore superior advantage in the analysis of WGS data on a large scale. The combination of DRAGEN™ and DeepVariant also provides a good balance of accuracy and efficiency as an alternative solution for germline variant detection in further applications. Our results facilitate the standardization of benchmarking analysis of bioinformatics pipelines for reliable variant detection, which is critical in genetics-based medical research and clinical application.

## Introduction

The innovation of next-generation sequencing (NGS) technologies has enabled exponential growth of the production of high throughput –omics data^1-3^. Whole genome sequencing (WGS) and targeted whole exome sequencing (WES) are two main types of DNA sequencing protocols that have been broadly applied for the discovery of disease-related genes and identification of driver mutations for specific disorders^4-6^. In contrast to WES, WGS can assess all the nucleotides of an individual genome and allow detection of variants in both coding and non-coding regions. As a result of decreasing genome sequencing cost, WGS is becoming a powerful tool to investigate a wide range of complex inherited genetic diseases (e.g. heart disease, diabetes and psychiatric conditions) through the identification of causal germline variants^7-11^. The clinical application of WGS is gaining utility and therefore importance in achievement of personalized precision medicine^12,13^.

There is a necessity of bioinformatic pipelines for variant calling analysis on WGS data in a precise and efficient way prior to its integration into clinical diagnostic application^14,15^. In general, a pipeline is comprised of the following steps: quality control, read alignment, variant calling, annotation, data visualization and reporting^12,16^. At the current stage of technological development, most of the clinical laboratories performing diagnostics of genetic disorders by WGS focus on two types of variants; single nucleotide polymorphisms (SNPs) and short insertion and deletions (indels). Many tools (e.g. Strelka, SpeedSeq, Samtools and Varscan2) have been developed for SNP and indel calling in the WGS analysis pipelines^17-20^. Among them, Genome Analysis Toolkit (GATK) is one of the most used variant calling tools, as it applies a variety of state-of-the-art statistical methods (e.g. logistic regression, Hidden Markov Chain and Naïve Bayes Classification) to accurately identify differences between the reads and the reference genome that are caused either by real genetic variants or by errors^21^. GATK can achieve high accuracy, but is still imperfect in memory management and running efficiency. Illumina® has released a Dynamic Read Analysis for GENomics (DRAGEN™) Bio-IT platform that provides an accurate and ultra-rapid solution for WGS data analysis^22^. The DRAGEN™ platform implements a highly configurable field-programmable gate arrays (FPGAs) hardware technique to dramatically speed up the analysis processes (e.g. alignment mapping and variant calling) and claims to do so without compromising accuracy. Verily Life Sciences (formerly Google Life Sciences) has developed the variant calling algorithm DeepVariant for small germline variant detection, based on a deep learning model^23^. DeepVariant applies the python TensorFlow library to call variants in aligned reads by learning statistical relationship between images of read pileups around putative variants and true genotype calls. In 2016, the PrecisionFDA Truth challenge reported DeepVariant as the most accurate pipeline in the performance of SNPs calling^24^.

To compare the accuracy and efficiency of different variant calling pipelines and score their competence, it is critical to have high-quality benchmark datasets in which the true variant calls are well known. The Genome in a Bottle Consortium (GiaB) developed a golden callset (sample NA12878) that is widely used during development of variant calling pipelines and benchmarking^25^. Since its release, the NA12878 callset has been continuously updated as a comprehensive resource, and one of the major improvements was integration of the truth callset independently generated by Platinum Genome (PG)^26^. An additional truth callset recently developed from a “synthetic-diploid” mixture of two haploid hydatidiform mole cell lines, CHM1 and CHM13, is now available in a public repository^27^. Although the variants in these two truth callsets represent real scenarios, the number of true variants is usually unknown, complicating its use for the assessment of accuracy (i.e. how close the defined truth callset is to the “true” mutational landscape). In contrast, simulated *in silico* WGS data allow users to generate variants under controlled scenarios with predefined parameters for which the “true” values are known, complementing the validation with real data. In several recent publications, performance comparisons of different pipelines, using both real and simulated WGS data, have shown to vary according to the dataset to which they have been applied^23,24,27-31^. Until now, none of the studies have evaluated the three pipelines (GATK, DRAGEN™ and DeepVariant) together using multiple sets of WGS data for benchmarking. Importantly, by combining different datasets, the accuracy of genomic variant identification can be compared in a more systematic way and a deeper understanding about their performance can be attained.

In this study, we obtained raw WGS data of NA12878 and “synthetic-diploid” samples from public repositories and constructed two sets of synthetic WGS data using read simulator. A comprehensive benchmarking of GATK, DRAGEN™, DeepVariant and their combinations was conducted using both real and simulated data. We aimed to evaluate the accuracy and efficiency of these pipelines for SNPs and short indels detection, to identify the most precise and efficient combination of tools for small variant calling. These were assessed according to performance, concordance and time consumption, in order to provide a useful guideline of reliable variant identification for genetic medical research and clinical application.

## Materials and Methods

### 1. Sources of WGS benchmarking dataset acquisition

#### 1.1. NA12878 (HG001) WGS data

The NIST reference material NA12878 (HG001) was sequenced at NIST, Gaithersburg, MD for the PrecisionFDA Truth Challenge. WGS library preparation was conducted using Illumina TruSeq (LT) DNA PCR-free sample Prep kits (FC-121-3001), and paired-end reads, insert Size: ∼550bp were generated on HiSeq 2500 platform with rapid run mode (2 flow cell per genome). Raw fastq files were obtained from https://precision.fda.gov/challenges/truth. In addition, another set of NA12878 raw WGS data sequenced in Supernat *et al.* was downloaded from the NCBI SAR repository (accession number: SRR6794144)^24^.

#### 1.2. “Synthetic-diploid” WGS data

Paired-end raw fastq files of “synthetic-diploid” WGS data were downloaded from the European Nucleotide Archive (accession number: SAMEA3911976). The reference material from a mixture of CHM1 (SAMN02743421) and CHM13 (SAMN03255769) cell lines at 1:1 ratio was sequenced on HiSeq X10 platform using PCR-free library protocol (Kapa Biosystems reagents)^27^. Two independently replicated runs, ERR1341793 and ERR1341796 are used for the benchmarking exercises.

#### 1.3. Simulated WGS data

In addition to real WGS data, reads were synthesized in silico using the tool Neat-GenReads^32^. Briefly, two independent sets of simulated paired-end reads in fastq format, together with true positive variant datasets in VCF format, were generated from a random mutation profile (average mutation rate: 0.002) and a userdefined mutation profile (using the golden truth callset assembled from CHM1 and CHM13 haploid cell lines, respectively). The simulation was performed on the basis of human reference genome build GRCH37 decoy, with a read length of 150 bp, an average coverage of 40X, and a median insert size of 350±70 bp.

### 2. Implementation of variant calling pipelines

Germline variant calling was performed using the pipelines: (1) GATK v4.1.0.0, (2) DRAGEN™ v3.7.0 and (3) DeepVariant v0.7.2 (see flowchart in Figure 1)^23,33^.

**Figure 1.**
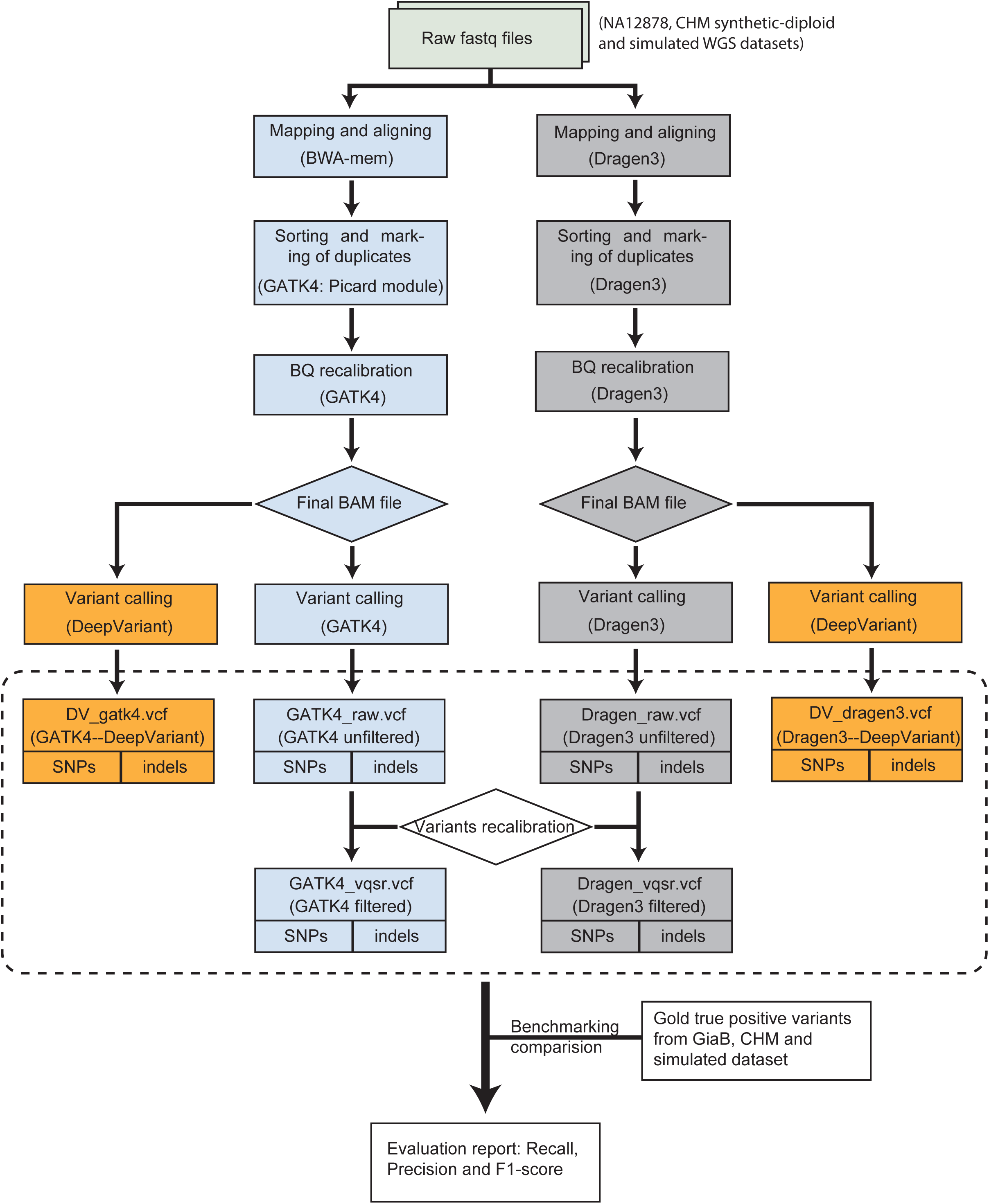
The flowchart of benchmarking analyses of different variant calling pipeline (GATK, DRAGEN™ and DeepVariant) combinations

**2.1.** The GATK pipeline workflow was applied following best practices (https://software.broadinstitute.org/gatk/best-practices). The raw paired-end reads were mapped to GRCH37.37d5 reference genome by BWA-mem v0.7.15^34^. Aligned reads were converted to BAM files and sorted based on the genome position after marking duplicates using Picard modules. The raw BAM files were refined by Base Quality Score Recalibration (BQSR) using default parameters. The variant calling (SNPs and indels) was performed by the HaplotyeCaller module. To speed up efficiency, the whole genome was split into 14 fractions and run in parallel, followed by merging of all runs into a final VCF file. Additionally, we used Variant Quality Score Recalibration (VQSR) to filter the original VCF files following GATK recommendations for parameter settings: HapMap 3.3, Omni 2.5, dbSNP 138, 1000 Genome phase I for SNPs training sets, and Mills- and 1000 Genome phase I data for indels.

**2.2.** The DRAGEN™ pipeline (https://www.illumina.com/products/by-type/informatics-products/dragen-bio-it-platform.html) followed a similar procedure as described for GATK best practices, including mapping and alignment, sorting, duplicate marking, haplotype calling and VQSR filtering.

**2.3.** The DeepVariant pipeline was run via singularity framework in accordance with online instructions (https://github.com/google/deepvariant/tree/r0.8/docs). In general, this consisted of three steps: (1) make_example module – consumes reads and the reference genome to create the TensorFlow example for evaluation with deep learning models. (2) call_variants module – consumes TFRecord files created by the make_example module and evaluates the model on each example in the input TFRecord. (3) postprocess_variants module – reads the output TFRecord files from the call_variants module, combines multi-allelic records and writes out a VCF file. DeepVariant only used transformed aligned sequencing reads for variant calling, and so processed BAM file from GATK or DRAGEN™ pipelines was fed as input.

Six VCF files were finally generated per each WGS dataset; there represent different parameter settings and processing combinations of the pipelines in terms of their workflows as depicted in Figure 1 (i.e. *DV_gatk4* – GATK for BAM file and DeepVariant for variant calling; *DV_dragen3* – DRAGEN™ for BAM file and DeepVariant for variant calling; *GATK4_raw* – GATK for both BAM file and variant calling; *GATK4_vqsr* – callset from *GATK4_raw* filtered with VQSR; *Dragen3_raw* – DRAGEN™ for both BAM file and variant calling and *Dragen3_vqsr* – callset from *Dragen3_raw* filtered with VQSR).

### 3. Computing environment and resources

Variant calling processes were run both on a high-performance computing (HPC) cluster and on a local virtual machine (VM) within the sensitive data platform (TSD) at the University of Oslo. The settings of each node in the HPC cluster include 64 AMD CPU cores with total 512 GB size of physical memory, the CentOS 7 operating system and BeeGFS network file system. The FPGA hardware infrastructure was installed on one node specific for DRAGEN™ pipeline application. The local VM has 40 CPU cores with a total 1.5 TiB physical memory, 2 TiB local disk with ext4 file system format and CentOS 7.

### 4. Benchmark consensus of VCF files

Gold standard truth callset with high confidence genomic intervals (NIST v3.3.2) for NA12878 (HG001) dataset were obtained from https://ftp-trace.ncbi.nlm.nih.gov/giab/ftp/release/NA12878_HG001/NISTv3.3.2/GRCh37/. To calculate the performance metrics, we used hap.py (version 0.3.8, vcfeval comparison engine) for comparison of diploid genotypes at haplotype level https://github.com/Illumina/hap.py. For benchmarking variants identified in simulated WGS data, we performed a consensus evaluation both with and without high-confidence regions. The variant callings of WGS data from the mixture of CHM1 and CHM13 were compared to the “synthetic-diploid” benchmark truth callset and high-confidence regions using the CHM-eval kit tool (version 20180222, vcfeval comparison engine)^27^. The definitions of true positive (TP), false positive (FP) and false negative (FN) were based on the types of variant matching stringencies “genotype match” (most strict) and “local match” (less strict)^35^. Precision, Recall and F1-score were calculated as TP/(TP+FP), TP/(TP+FN) and 2*TP/(2*TP+FN+FP), respectively.

## Results

### 1. Quality summary of WGS datasets

Two NA12878 WGS datasets, derived from PrecisionFDA and SRR679144, have 542,906,383 and 379,033,340 read pairs with a median insert size 553 bp and 540 bp, and average coverage ∼50X and ∼37X (Table S1). For “synthetic-diploid” datasets, two independent replicate runs have 414,011,224 and 514,732,237 read pairs, with a median insert size 354 bp and 329 bp, and a sequencing depth of ∼40X and ∼50X, respectively. About 98.7% - 99.4% of sequencing reads in real WGS datasets could be aligned to the reference genome (GRCh37.hs37d5). In comparison, two simulated datasets, Sim_random and Sim_user, have 390,319,108 and 390,296,059 read pairs with a sequencing depth of ∼40X, and almost 100% of reads could be aligned to the reference genome (Table S1). Of all datasets, NA12878_SRR679144 displays an unexpectedly high duplicate level of mapped reads (26%) compared to the others (0.2%-2.6%). We called variants for both real and simulated data with the defined bioinformatic pipelines (Figure 1), which resulted in six VCF files (see details in Materials and Methods) per each dataset.

### 2. Benchmarking of GATK, DRAGEN and DeepVariant variant calling pipelines

The accuracy of germline variant calls using NA12878 and “synthetic-diploid” WGS datasets was compared firstly. For SNP calls, all benchmarked pipelines (and their combinations) have F1-score, recall and precision values higher than 0.963, 0.932 and 0.986 respectively. Specifically, *Dragen3_raw* shows the highest F1-score value in NA12878_PrecisionFDA dataset, while *DV_Dragen3* outperforms others in F1-score for NA12878_SRR6794144 dataset (Figure 2A and 2C). *DV_gakt4* has the best performance in accuracy for the two replicate runs of “synthetic-diploid” datasets (Figure 2E and 2G). Furthermore, we found that F1-scores in five of the six combinations are closer to each other, except for *GATK4_vqsr* with a range of values 0.989 - 0.996. The lower F1-score of *GATK4_vqsr* is mainly due to poor performance in recall metrics, although precision metrics can reach a high value in real datasets (Figure 2).

**Figure 2.**
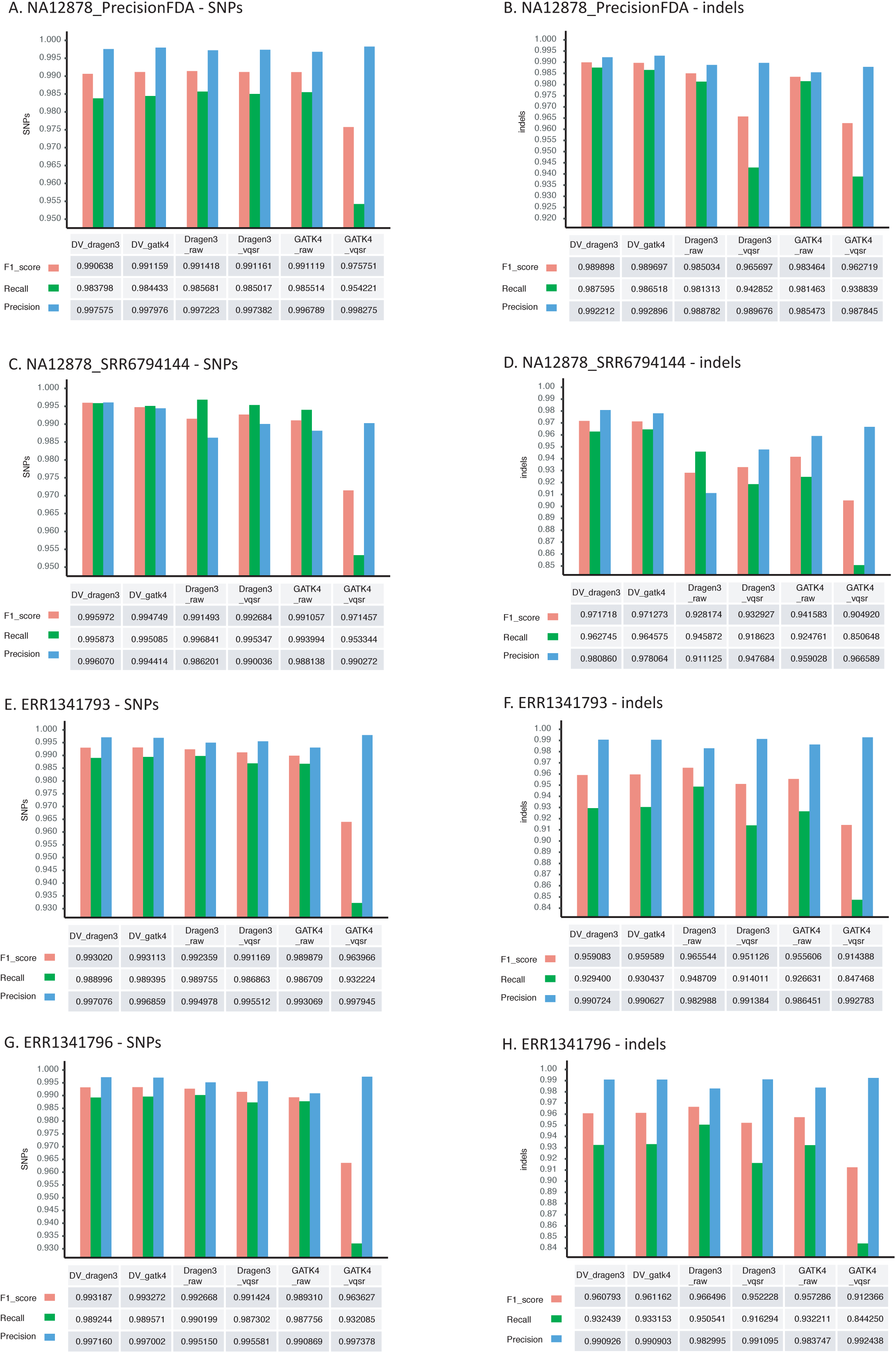
Accuracy evaluation of variant calling pipelines on real WGS datasets NA12878_PrecisionFDA (A and B), NA12878_SRR6794144 (C and D) and “synthetic-diploid” CHM1-13 (E and F for replicate ERR1341793, G and H for replicate ERR1341796). For each dataset, six different combinations (i.e. *DV_gatk4, DV_dragen3, Dragen3_raw, Dragen3_vqsr, GATK4_raw* and *GATK4_vqsr*) were compared. The performance metrics (F1-score, Recall and Precision) of SNP and indel calls were estimated using “genotype match” approach for NA12878 and “local match” approach for “synthetic-diploid” CHM1-13.

Compared to SNP calls, the performance of indel calls is more diverse; F1-scores range from 0.905 to 0.989 in NA12878 dataset and from 0.912 to 0.961 in the “synthetic-diploid” dataset (Figure 2B, 2D, 2F and 2H). Notably, *DV_dragen3* shows a higher F1-score than others in two datasets of NA12878, whereas the accuracy of *Dragen3_raw* gives the best performance in two replicate runs of “synthetic-diploid”. Again, *GATK4_vqsr* has a poor performance in F1-score regardless of datasets. By contrast, the benchmark evaluation on two simulated WGS datasets shows similar F1-score metrics for SNP and indel calls, respectively, in which *Dragen3_raw* gives the best performance in accuracy regardless of whether the benchmarking is done with a high confidence bed file or not (Figure S1). In total, our results indicate *Dragen3_raw* and *DV_dragen3* achieve a better F1-score for small variant calls in the analyses of real and simulated datasets.

### 3. Stratification analysis of different genome contexts

We stratified the performance and evaluated benchmarking metrics in different genome contexts. Recall, precision and F1-scores that were compared in conserved and coding regions for “synthetic-diploid” datasets are displayed in Table S2. The performances of all pipeline combinations (except *GATK4_vqsr*) are similar to each other, with F1-score ranging from 0.9944 to 0.9967 for SNP calls. Although metrics of indel calls are variable in F1-score, differences between *DV_gatk4, DV_dragen3, Dragen3_raw* and *GATK4_raw* are not significant (Table S2). Similarly, stratification analysis on conserved/coding regions using simulated WGS datasets shows analogous F1-scores among different pipeline combinations (Table S3). In addition, the performance in low complexity regions and regions with various GC content was evaluated. As expected, the metric values (F1-score, recall and precision) of SNPs and indels tend to decrease with the increase in abundance of tandem repeats, and all pipelines give a poor accuracy of variant calling in the low complexity regions with repeat length > 200 bp (Figure S2). GC stratification analysis on SNPs and indels shows a similar pattern, with a poor performance of F1-score in regions of high and low GC content (Figure S3).

The analysis of the substitution signature and context of false positive and negative variants in NA12878_SRR6794144 dataset demonstrated that there are more calls with A>T, C>A, G>T and T>A substitutions in *GATK4_raw* false positive variants than expected from the distribution of variants in the true gold callset (Figure S4A), which supports earlier findings reported by Supernat *et al.* Additionally, more C>A substitutions in both false positive and negative variants called by *Dragen3_raw* were found. In comparison to NA12878_PrecisionFDA dataset, more A>C and T>G substitutions were identified in false positive variants called by *GATK4_raw*, than was expected (Figure S4B). However, in the simulated datasets, both false positives and negatives called by all pipeline combinations seem to be independent of compositional biases with respect to the base change (Figure S4C and S4D).

### 4. Comparison of variant calling concordance

Venn diagrams in Figure 3 illustrate the intersection of SNPs and indels called by *GATK4_raw, Dragen3_raw, DV_gatk4* and *DV_dragen3*. For both real and simulated datasets, around 91.7%-99.6% SNPs were jointly reported by all the pipeline combinations, and over 95.3%-99.95% of SNPs can be detected by at least two pipeline combinations. The fractions of SNPs uniquely called by *GATK4_raw, Dragen3_raw, DV_gatk4* and *DV_dragen3* were only 0.002%-1.62%, 0.045%-2.56%, 0.005%-0.31% and 0.0004%-0.32%, respectively. By contrast, there are 83.5%-99.4% indel variants commonly detected by multiple combinations in all datasets except for NA12878_SRR6794144, in which only 69.7% of total indels were jointly identified (Figure 3). Although indels have a larger divergence of calling concordance compared to SNPs, the high number of variants detected by multiple combinations and low orphan variants support a good agreement in the identification of SNPs and indels by different pipelines.

**Figure 3.**
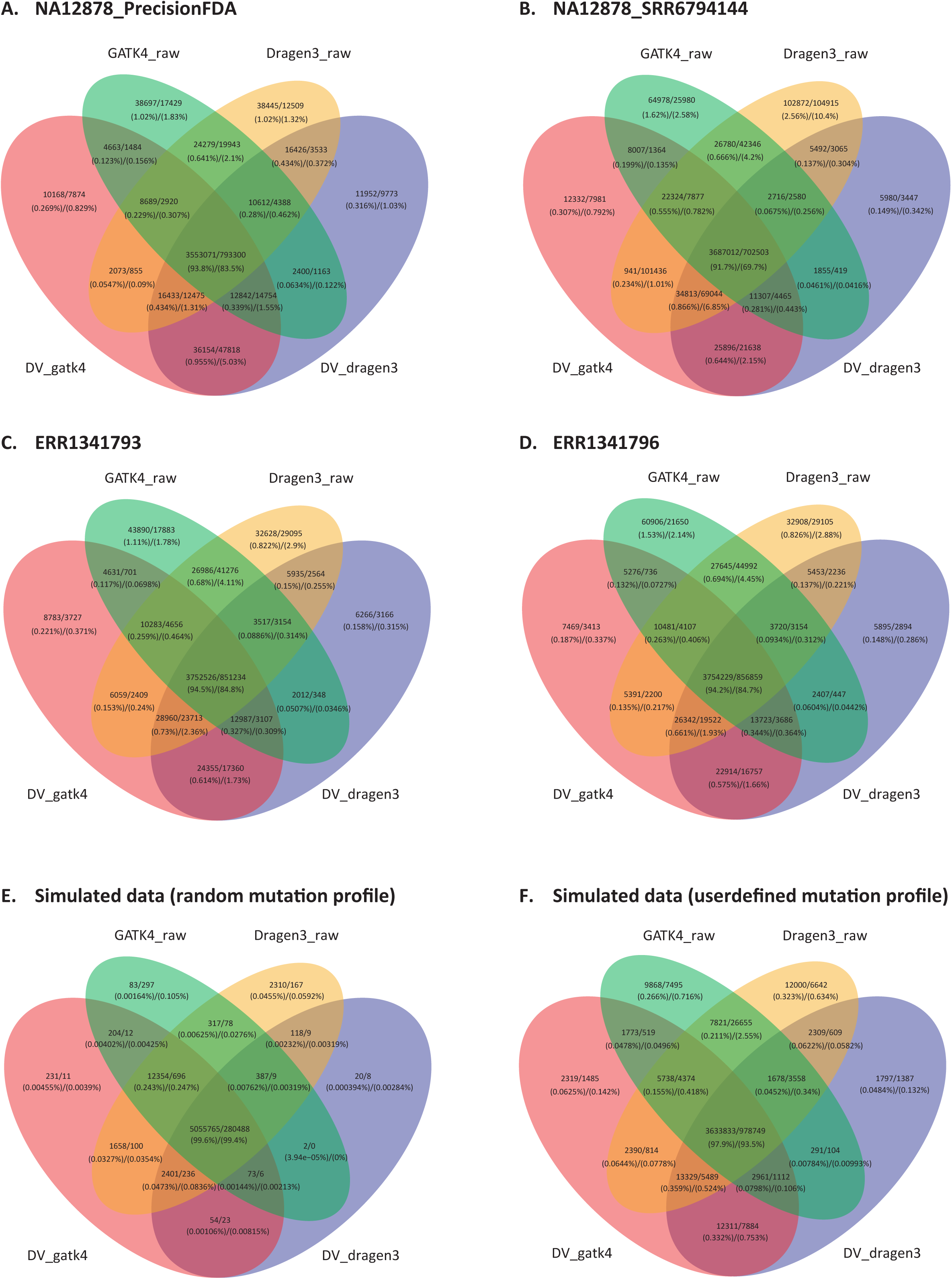
Venn diagrams showing the intersection of variants called by different pipeline combinations on NA12878_PrecisionFDA (A), NA12878_SRR6794144 (B), “synthetic-diploid” CHM1-13 (C – replicate ERR1341793; D – replicate ERR1341796) and two simulated WGS datasets (E – random mutation profile; F – userdefined mutation profile). The number of SNP and indel variants are shown together using separator ‘/’. The callsets *Dragen3_vqsr* and *GATK4_vqsr* are not included in comparison as they are a subset of *Dragen3_raw* and *GATK4_raw*, respectively.

### 5. Comparison of execution time

To better assess the operating efficiency, the pipeline processing procedure was divided into upstream (Fastq to BAM file) and downstream (BAM to VCF file) workflows, and runtime of each workflow was measured. For benchmarking execution time on HPC cluster, *Dragen3_raw/vqsr* took from 30 minutes to 1.5 hours in the upstream analysis. This was significantly lower than *GATK4_raw/vqsr*, with a speed-up gain in the range of 17X to 33X (Figure 4). In the downstream workflow, *Dragen3_raw/vqsr* still outperforms *GATK4_raw/vqsr* and *DV_gatk4/DV_dragen3*, despite the degree of speed-up gain being less than that in the upstream workflow. Similarly, DRAGEN™ shows a big advantage in running speed when a comparison was done on a local VM, with a time requirement of even less than benchmarking on HPC cluster (Figure S5). Overall, compared to the other pipelines, DRAGEN™ platform provides an ultra-rapid analysis solution for germline variant calling using WGS data.

**Figure 4.**
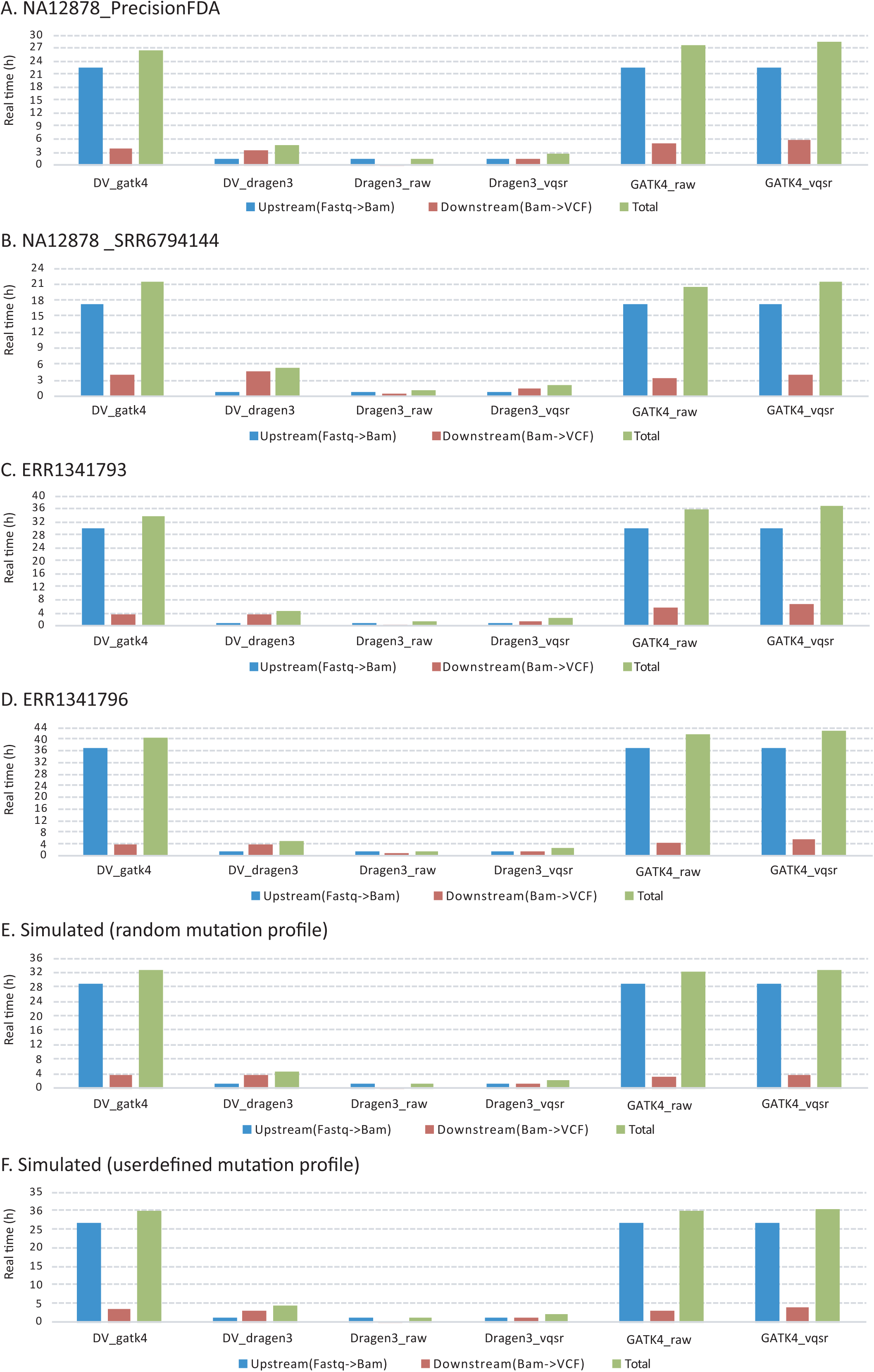
Variant calling runtime of six pipeline combinations (*DV_gatk4, DV_dragen3, Dragen3_raw, Dragen3_vqsr, GATK4_raw* and *GATK4_vqsr*) benchmarked on HPC cluster (A and B – NA12878_PrecisionFDA and NA12878_SRR6794144 datasets; C and D – “synthetic-diploid” ERR1341793 and ERR1341796 datasets; E and F - simulated data based on a random and a userdefined mutation profiles).

## Discussion

In this study, we empirically evaluated the performance of different pipelines (and their combinations) for germline variant calling using real and simulated WGS data. Our results demonstrate that DeepVariant (*DV_dragen3* or *DV_gatk4*) shows a higher accuracy in SNP calls for one NA12878 dataset (SRR6794144) and two “synthetic-diploid” datasets, and in indel calls for two NA12878 datasets. Despite a better performance, the F1-scores obtained in NA12878 benchmarking evaluation are lower than that published in the FDA Truth Challenge: 0.9912-0.9959 versus 0.9996 (pFDA top) for SNP calls, and 0.9897-0.9717 versus 0.9934 (pFDA top) for indel calls. This variation probably results from differences in the benchmarking procedure of pFDA Truth challenge, in which the NA12878 sample was used for training and the HG002 sample was used for testing. The top benchmarking results in pFDA Truth challenge are derived from the HG002 comparison. The accuracy of the DRAGEN™ pipeline (*Dragen3_raw*) gave a better performance in both SNP and indel calls for the simulated dataset, and in indel calls for the “synthetic-diploid” datasets, despite not achieving as high F1-score metrics as DeepVariant in the benchmark of the NA12878 dataset. In fact, the differences in benchmarking scores between DRAGEN™ and DeepVariant are quite small (Figure 2 and Figure S1). In particular, stratification analysis of conserved and coding regions suggests nearly the same accuracy between them.

As the DRAGEN™ platform was developed for speed, accuracy, flexibility and cost effectiveness, its most important advantage is computational time and throughput of the massive volumes of data. Indeed, in this study the running efficiency of the DRAGEN™ platform was far superior to GATK and DeepVariant. Based on these considerations, and the accuracy results we measured, it is recommended that either the DRAGEN™ pipeline is used alone (*Dragen3_raw*) or in combination (*DV_Dragen3*), where DRAGEN™ is used for upstream processing and DeepVariant for downstream processing, to provide a balance in accuracy and efficiency for germline variant calling from WGS data.

Although the DRAGEN™ platform gave the best performance in running efficiency in this study, real execution time on HPC clusters never reached the performance stated by the manufacturer. Even when a benchmarking comparison was performed on a local VM, where faster I/O communication on the local ext4 file system can benefit the running speed compared to BeeGFS network file system on a HPC cluster, only a minor improvement in time consumption was observed (Figure S5). Thus, there is still room for optimizing the runtime of DRAGEN™ platform with regards to its implementation at e-frastructure and hardware level. In comparison to DRAGEN™, the optimization of running efficiency for GATK and DeepVariant was not achieved in the computing environment of our study. For example, DeepVariant could gain 2.5X speedup using a high-performance graphics processing unit, since its variant calling algorithm is based on image analyses. For GATK, the genome was split into 14 fractions by chromosomes, scaffolds and contigs, and were run in a “scatter-gather” strategy. There are 64 cores per node in the HPC cluster, therefore the genome could ideally be split into the same number of divisions as the number of cores, and be run in parallel. Despite these optimizations, neither DeepVariant nor GATK will achieve the efficiency of DRAGEN™, as no hardware-accelerated implementations of genomic analyses algorithms have been developed for them.

Two types of high confidence benchmark truth call sets: the GiaB reference data (sample NA12878) and the “synthetic-diploid” mixture of two haploid cell lines were applied to evaluate the performance of germline variant calls using real WGS data. The construction of the truth set, and strengths and weaknesses based on variant type and genome context should be considered. The GiaB benchmark sets were built from the consensus of multiple variant callers on Illumina short-read sequencing with the aid of a pedigree analysis, integration of structural variants identified with long fragment technologies by PacBio and 10X Genomics, and HuRef genome analysis using Sanger sequencing^35^. Nearly all the “true” variants in NA12878 sample are present in the resource files (e.g. dbSNP, 1000 genomes and the training data for DeepVariant) used for pipeline running. In this case, the results are likely overfit as the answer has been used all along. Furthermore, NA12878 tends to exclude more difficult types of variants in the region with moderately diverged repeats, and segmental duplications, as consensus in such regions has not been reached. This will tend to bias GiaB datasets towards “easy-to-sequence-and-analyze” genome regions.

The truth “synthetic-diploid” callset was generated by assembly of long reads sequenced from two haploid cell lines (CHM1 and CHM13) using PacBio technology. This can be considered trustworthy as there are no heterozygous sites that tend to confuse the assembly. The exclusive use of PacBio, without incorporation of the flaws generated from Illumina’s short-read technology, ensure there is less correlation between the failure modes of this method on the short-read data and confidence regions. This enables benchmarking in regions that are difficult to map with short reads. However, the “synthetic-diploid” call set currently contains some errors that were intrinsically present in the long reads^27^. It is thus recommended to use a less strict benchmarking strategy (“local matches” method) for comparisons^27,35^. Here, the evaluation using “genotype match” as it applied in NA12878 datasets was performed as well (Table S4). For SNP metrics, DeepVariant (*DV_gatk4* or *DV_dragen3*) was consistently rated as the best according to their respective F1-scores. In terms of the performance metrics of indel calls, *Dragen3_raw* and *GATK4_raw* have a better value for ERR1341793 and ERR1341796 datasets, respectively. As expected, the recall, precision and F1-score of indels are relatively low compared to the metrics done by the “local match” method. Precisely assessing the accuracy of genotypes from the exact sequence changes in the REF and ALT fields of the VCF file for “synthetic-diploid” data benchmarking remains challenging. Consequently, a less stringent methodology like the “local match” approach is required. One advantage is robust towards representational differences of variants in truth and inquiry sets. Overall, the characteristics of these two truth datasets make them invaluable for performing a comprehensive comparison assessment of different bioinformatics tools.

In addition to the real WGS data, we generated two simulated WGS datasets on the basis of a random and a userdefined mutation profile. One advantage of using simulated *in silico* data for benchmarking is that all “true” positive SNPs and indels are known. The calculation of F1-score is more accurate due to the reduced risk of overestimating false negatives. Additionally, in simulated data the read coverage across the whole genome region has a more even distribution than that found in real data, so variant calling errors arising from low coverage in some regions could be reduced. Moreover, read simulation is able to avoid false positive calls naturally originating from random or systematic noise in the real data. As seen in Figure S4, false positive calls in simulated datasets are independent of compositional biases in the distribution of substitution signatures. On the other hand, the accuracy of variant calling in simulated data easily reaches saturation (Figure S1), as simulated WGS data can achieve a perfect alignment (almost 100%) to the reference genome (Table S1). This is because the models used for data simulation may not replicate the sequence complexity present in real data precisely. For examples, some important modelling parameters such as PCR amplification during library preparation, GC% coverage bias, sequencing errors and mutation profile were empirically learned from selected known datasets without considering sample specificity broadly. Although the models do not fit real scenarios of genome context and structure completely, simulation alone is still an important approach for benchmarking evaluation of different pipelines with similar functionality. It can allow users to validate biological models and gain understanding about specific dataset, which is helpful for the development of new computational tools.

It is highly recommended in GATK and DRAGEN™ best practices to apply variant quality score recalibration (VQSR) to filter raw SNP and indel calls generated by HaplotypeCaller, and to remove calling artefacts. In theory, VQSR balances sensitivity and specificity during variant filtering. However, the F1-score was lower in both real and simulated data except for *Dragen3_vqsr* in NA12878_SRR679414 after VQSR filtering, although precision reached the highest value. In Figure 2, the precision metrics on average were raised only 0.15% and 0.5% for SNPs and indels, respectively, but the recall suffered from a larger fall, which is significant for *GATK4_vqsr* (e.g. reduced by 3% for SNPs and 4% indels in NA12878_PrecisionFDA dataset). Consequently, the calculated F1-score did not give the expected improvement. Such result could be potentially explained by the fact that VQSR was performed on a single sample at a time, yielding instability from the convergence failure of core algorithm modelling. This may lead to the necessity for quite “strict” criteria in the filtering of raw variant calls and cause a lower recall value. In addition, we experienced some challenges in performing VQSR analysis on the simulated WGS data under the default parameters, as there were not enough variants to be trained as a meaningful “bad set”, which is used for cluster discrimination. Instead, we turned down the number of max-gaussian parameters to 2 for indels and 4 for SNPs and forced the program to group variants into a smaller number of clusters to satisfy the statistical requirements. Overall, our results suggest there is no necessity to perform VQSR control for one sample analysis, and in fact the raw unfiltered VCF files have a good balance between recall and precision for GATK and DRAGEN™. GATK developers are currently working on a new deep learning-based filtering method to classify and refine variants, which could potentially overcome the weakness of VQSR in further work.

In conclusion, our benchmarking exercises on real and simulated WGS datasets reveal DRAGEN™ and DeepVariant pipelines have a high accuracy in small germline variant calling, and there are no significant differences in their F1-score performances. However, the DRAGEN™ platform performed superiorly in ultra-rapid analysis of WGS data for SNPs and indel detection, and therefore has great potential for implementation in routine genomic medicine. The combination of DeepVariant and DRAGEN™ pipelines can also offer a fast, efficient and reliable way to analyze WGS data on a large scale, and go a long way toward reliable and consistent calling of variants when translating genetic variant information to medical diagnoses.

## Data availability

The synthetic whole genome sequencing data generated and analyzed in this study are available from the corresponding authors on request.

## Acknowledgement

We thank Ghislain Fournous for suggestions on parameter settings of GATK pipelines, Olav Pedersen for his contribution in performing benchmarking and Serena Elizabeth Marshall for her contribution in reviewing the manuscript. The study was funded by grants from the Research Council of Norway through BigMed project. We also acknowledge Services for Sensitive Data (TSD) at the University of Oslo for secure storage of data and high-performance computing support (project number: p21).

## Author contributions

Study concept and design: E.H, S.Z, O.A; Acquisition and collection of data: S.Z, O.A and T.S; Implementation of tools, data analysis and interpretation: S.Z, O.A and A.A; Drafting of the manuscript: S.Z; Critical revisions and comments of the manuscript: S.Z, O.A, T.S, A.A, E.H; Funding providing and study supervision: E.H

