## Supplementary figure for "Accuracy and efficiency of germline variant calling pipelines for human genome data"

A. Simulated (random mutation profile) - SNPs

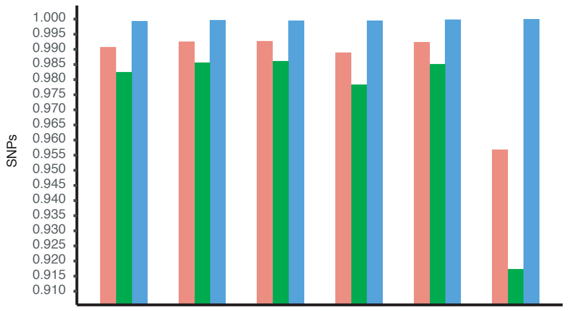

| | DV_dragen3 | DV_gatk4 | Dragen3_raw | Dragen3_vqsr | GATK4_raw | GATK4_vqsr |
|---|---|---|---|---|---|---|
| F1_score | 0.990803 | 0.992578 | 0.992786 | 0.988857 | 0.992401 | 0.956906 |
| Recall | 0.982421 | 0.985613 | 0.986086 | 0.978365 | 0.985109 | 0.917378 |
| Precision | 0.999330 | 0.999643 | 0.999579 | 0.999576 | 0.999802 | 0.999993 |

B. Simulated (random mutation profile) - indels

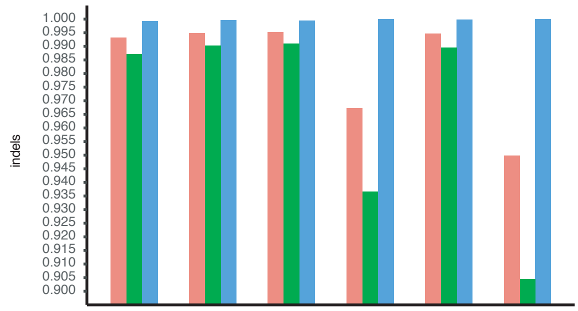

| | DV_dragen3 | DV_gatk4 | Dragen3_raw | Dragen3_vqsr | GATK4_raw | GATK4_vqsr |
|---|---|---|---|---|---|---|
| F1_score | 0.993141 | 0.994948 | 0.995262 | 0.967213 | 0.994646 | 0.949805 |
| Recall | 0.987064 | 0.980329 | 0.991032 | 0.936558 | 0.989553 | 0.904424 |
| Precision | 0.999293 | 0.999610 | 0.999529 | 0.999944 | 0.999791 | 0.999981 |

C. Simulated (userdefined mutation profile) - SNPs

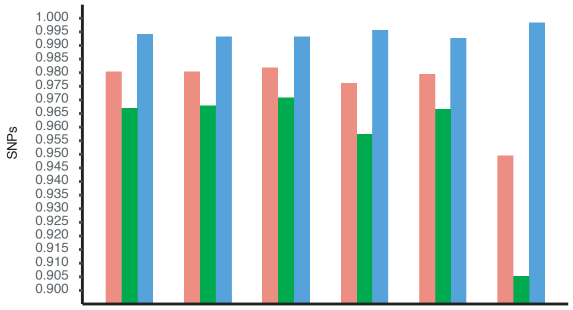

| | DV_dragen3 | DV_gatk4 | Dragen3_raw | Dragen3_vqsr | GATK4_raw | GATK4_vqsr |
|---|---|---|---|---|---|---|
| F1_score | 0.980336 | 0.980320 | 0.981865 | 0.976108 | 0.979361 | 0.949287 |
| Recall | 0.966984 | 0.967792 | 0.970702 | 0.957356 | 0.966482 | 0.905098 |
| Precision | 0.994062 | 0.993177 | 0.993289 | 0.995611 | 0.992588 | 0.998013 |

D. Simulated (userdefined mutation profile) - indels

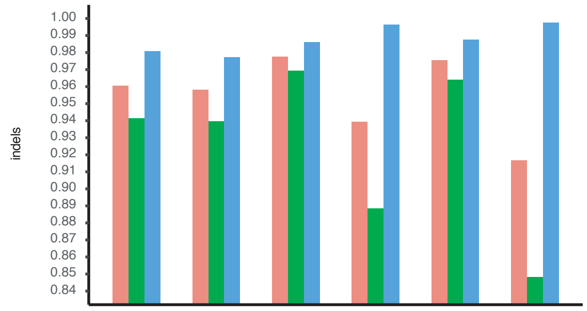

| | DV_dragen3 | DV_gatk4 | Dragen3_raw | Dragen3_vqsr | GATK4_raw | GATK4_vqsr |
|---|---|---|---|---|---|---|
| F1_score | 0.960696 | 0.958139 | 0.977657 | 0.939439 | 0.975587 | 0.915420 |
| Recall | 0.941496 | 0.939597 | 0.969263 | 0.888650 | 0.963996 | 0.847836 |
| Precision | 0.980696 | 0.977428 | 0.986197 | 0.996385 | 0.987460 | 0.994711 |

E. Simulated (random mutation profile) - SNPs

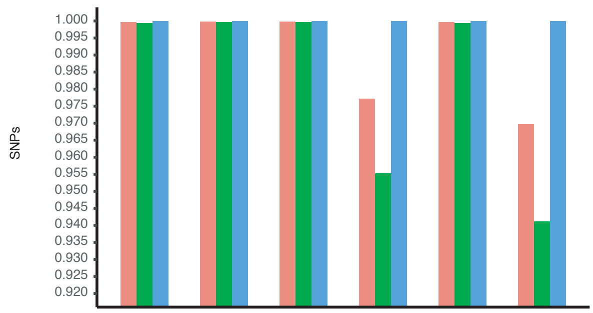

| | DV_dragen3 | DV_gatk4 | Dragen3_raw | Dragen3_vqsr | GATK4_raw | GATK4_vqsr |
|---|---|---|---|---|---|---|
| F1_score | 0.999793 | 0.999626 | 0.999836 | 0.977101 | 0.999688 | 0.969658 |
| Recall | 0.999622 | 0.999322 | 0.999712 | 0.955231 | 0.999392 | 0.941118 |
| Precision | 0.999965 | 0.999930 | 0.999961 | 0.999996 | 0.999984 | 0.999983 |

F. Simulated (random mutation profile) - indels

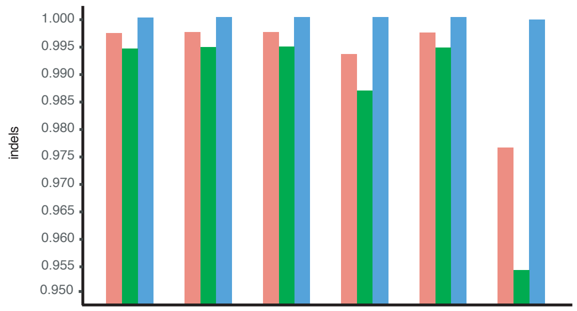

| | DV_dragen3 | DV_gatk4 | Dragen3_raw | Dragen3_vqsr | GATK4_raw | GATK4_vqsr |
|---|---|---|---|---|---|---|
| F1_score | 0.997224 | 0.997089 | 0.997290 | 0.993231 | 0.997193 | 0.976691 |
| Recall | 0.994495 | 0.994258 | 0.994618 | 0.986578 | 0.994429 | 0.954448 |
| Precision | 0.999968 | 0.999935 | 0.999975 | 0.999975 | 0.999973 | 0.999996 |

G. Simulated (userdefined mutation profile) - SNPs

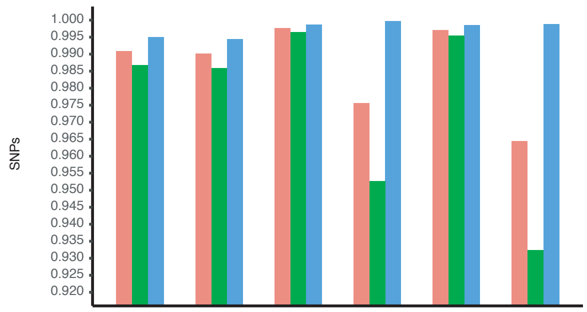

| | DV_dragen3 | DV_gatk4 | Dragen3_raw | Dragen3_vqsr | GATK4_raw | GATK4_vqsr |
|---|---|---|---|---|---|---|
| F1_score | 0.990099 | 0.990898 | 0.997584 | 0.975621 | 0.997001 | 0.964460 |
| Recall | 0.985867 | 0.986770 | 0.996433 | 0.952716 | 0.995449 | 0.932326 |
| Precision | 0.994368 | 0.995060 | 0.998739 | 0.999654 | 0.998557 | 0.998889 |

H. Simulated (userdefined mutation profile) - indels

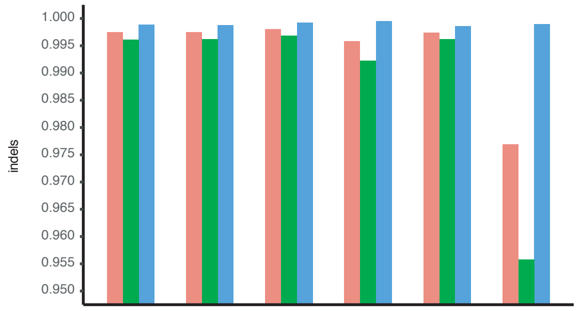

| | DV_dragen3 | DV_gatk4 | Dragen3_raw | Dragen3_vqsr | GATK4_raw | GATK4_vqsr |
|---|---|---|---|---|---|---|
| F1_score | 0.997502 | 0.997477 | 0.998048 | 0.995880 | 0.997395 | 0.976920 |
| Recall | 0.996194 | 0.996109 | 0.996853 | 0.992246 | 0.996226 | 0.955811 |
| Precision | 0.998814 | 0.998849 | 0.999245 | 0.999541 | 0.998566 | 0.998983 |

**Figure S1.** Accuracy evaluation of variant calling pipelines on two ”simulated” WGS datasets generated using random mutation profile (A and B - without using high conf. region bed file; E and F - using high conf. region bed file) or userdefined mutation profile (C and D - without using high conf. region bed file; G and H - using high conf. region bed file), respectively. For each dataset, six different combinations (DV_gatk4, DV_dragen3, Dragen3_raw, Dragen3_vqsr, GATK4_raw and GATK4_vqsr) were compared. The performance metrics (F1-score, Recall and Precision) of SNP and indels calls were estimated using “genotype match” approach.

**Figure S2A.** Stratification analysis of SNP callings in low complexity regions (benchmarked without using high conf. regions)

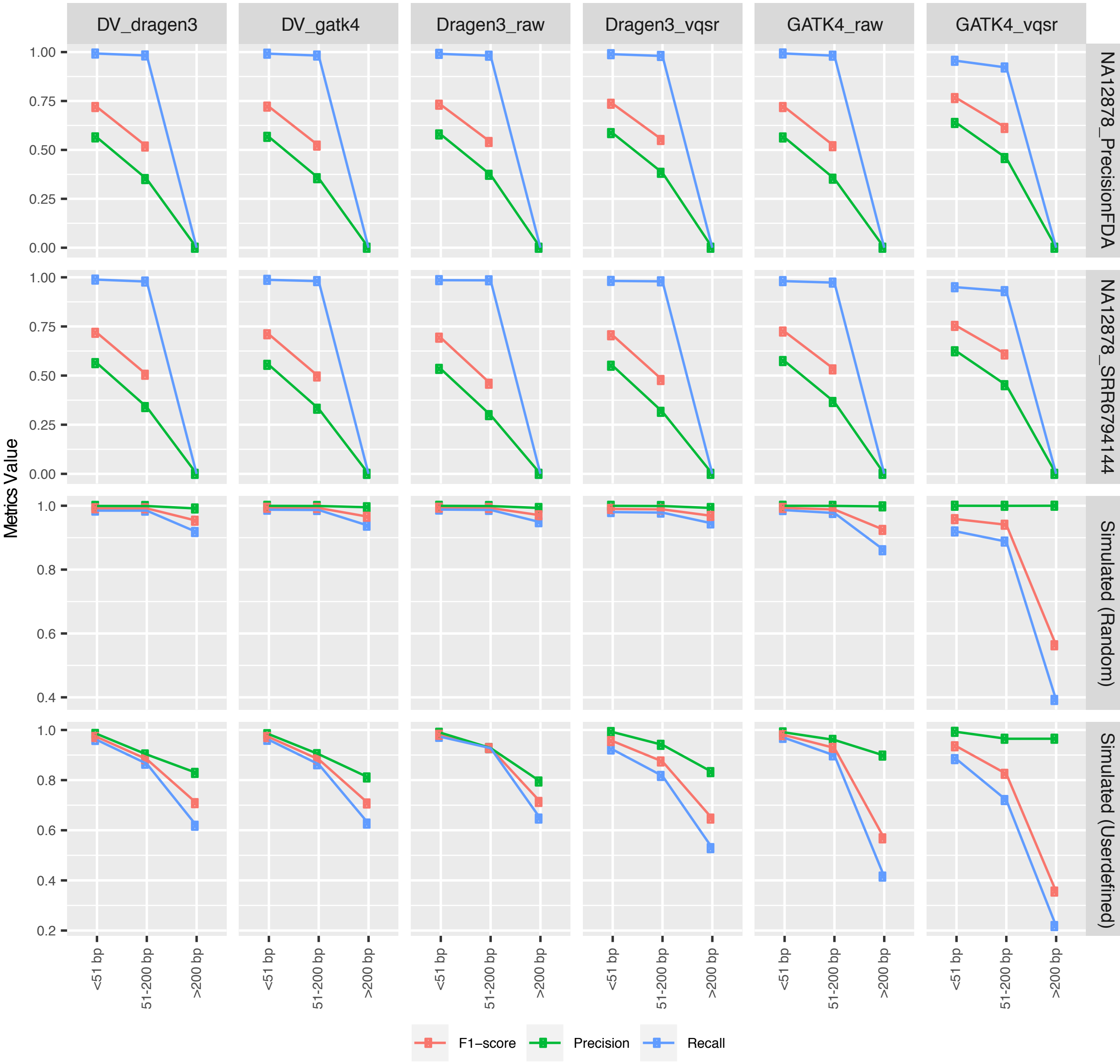

**Figure S2B.** Stratification analysis of indel callings in low complexity regions (benchmarked without using high conf. regions)

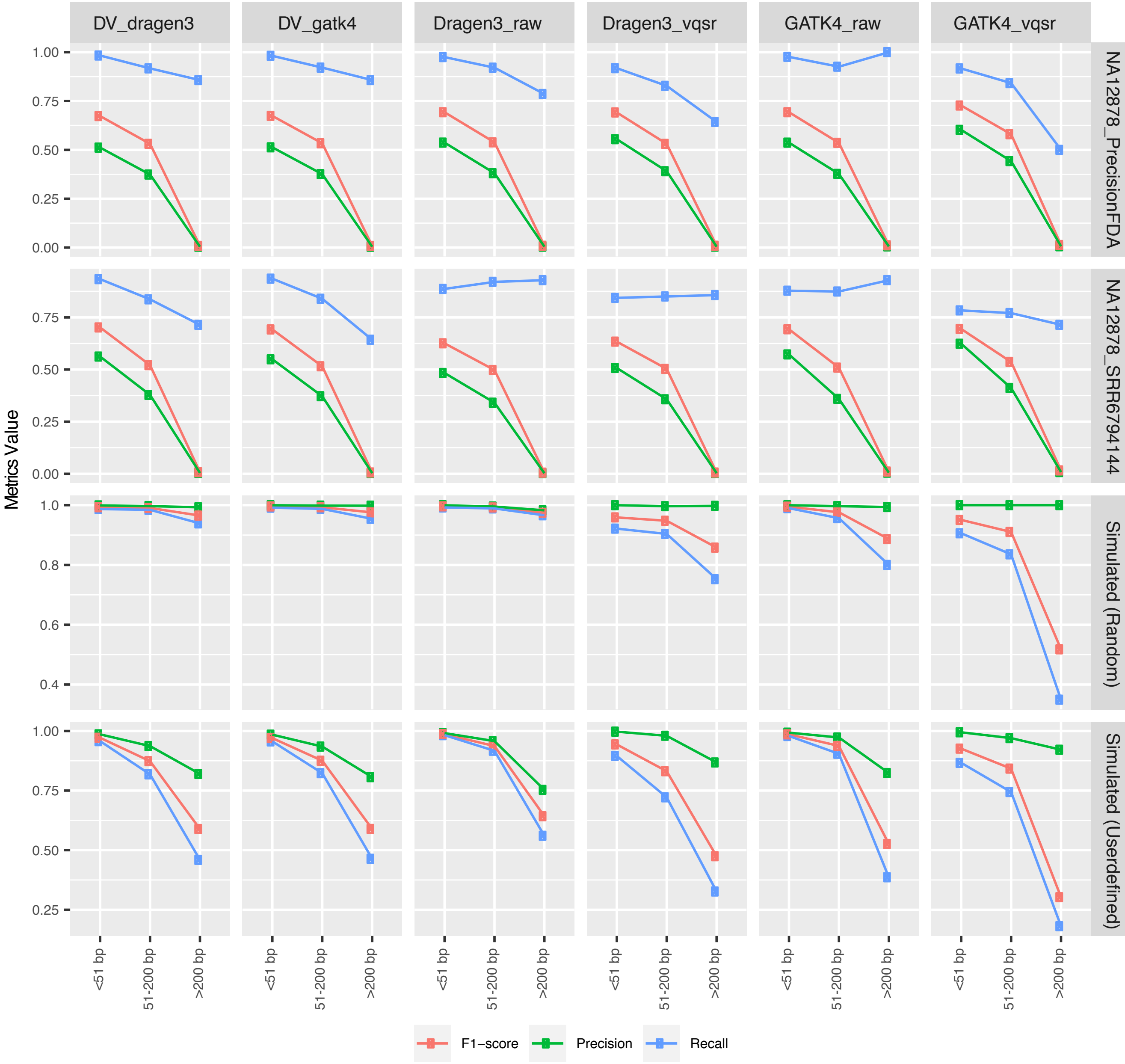

**Figure S3A.** Stratification analysis of SNP callings in different GC content regions (benchmarked without using high conf. regions)

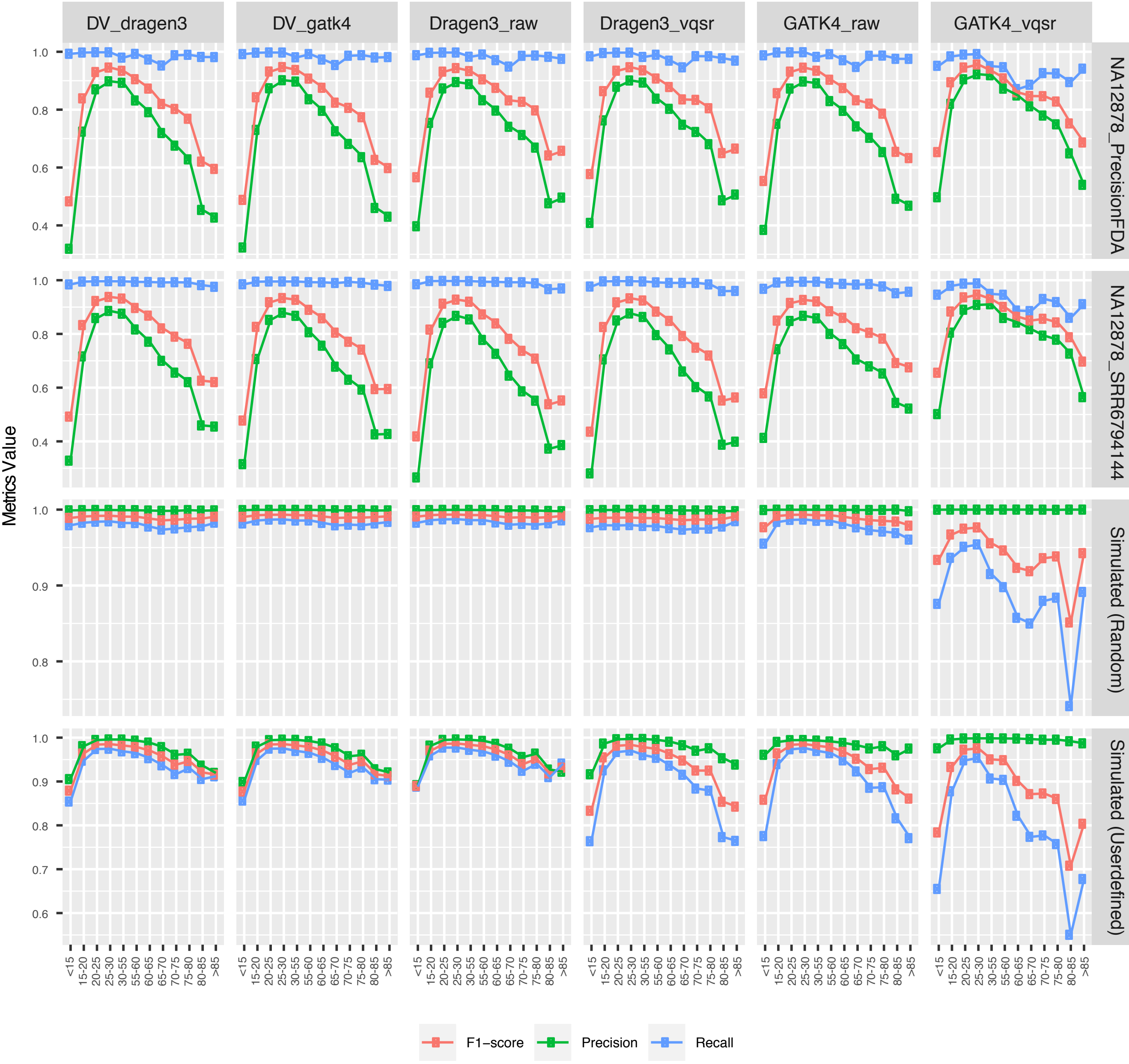

**Figure S3B.** Stratification analysis of indel callings in different GC content regions (benchmarked without using high conf. regions)

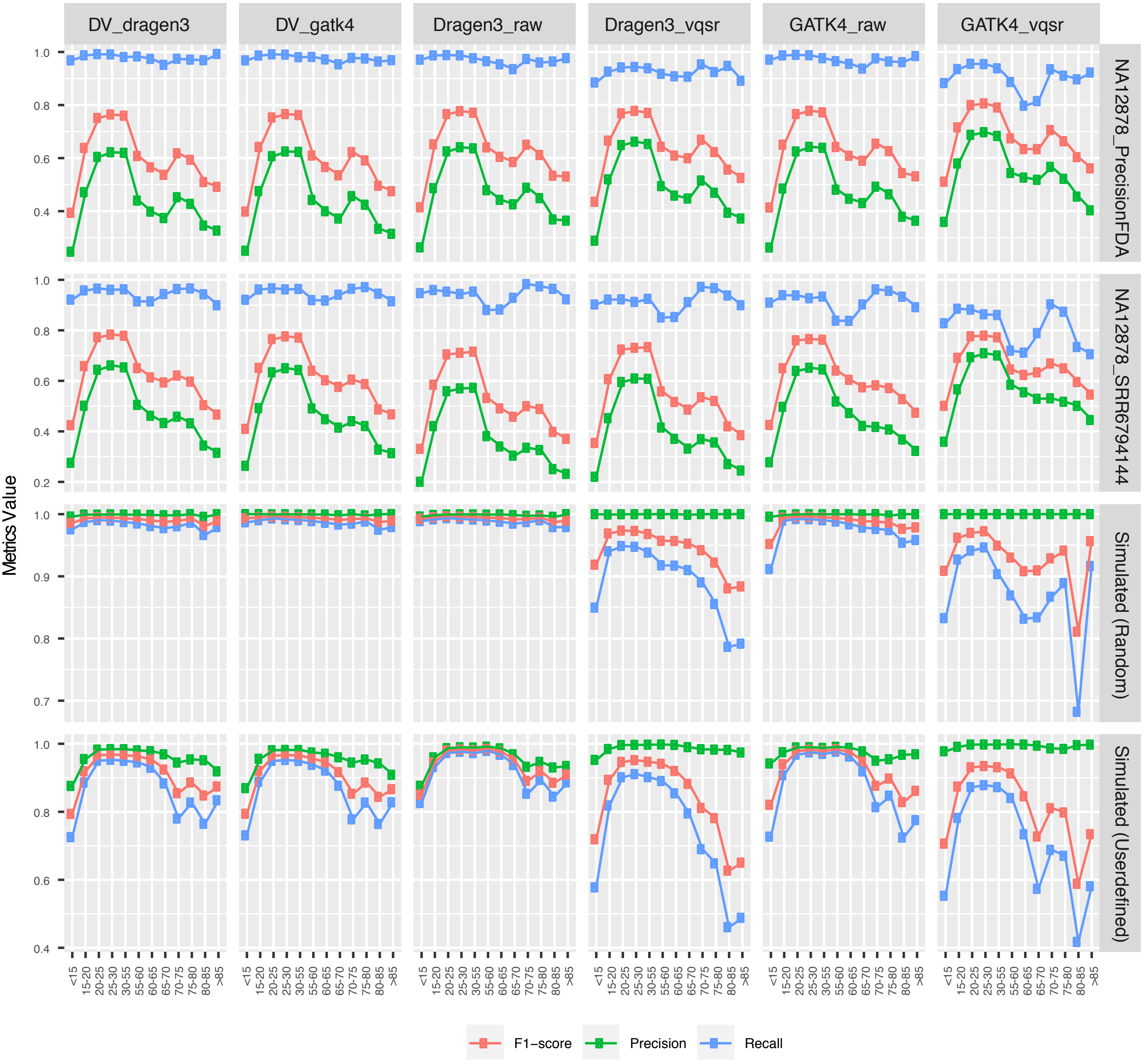

**Figure S4A.** The distribution of substitution signature of false positive and negative variants for NA12878_SRR6794144 dataset

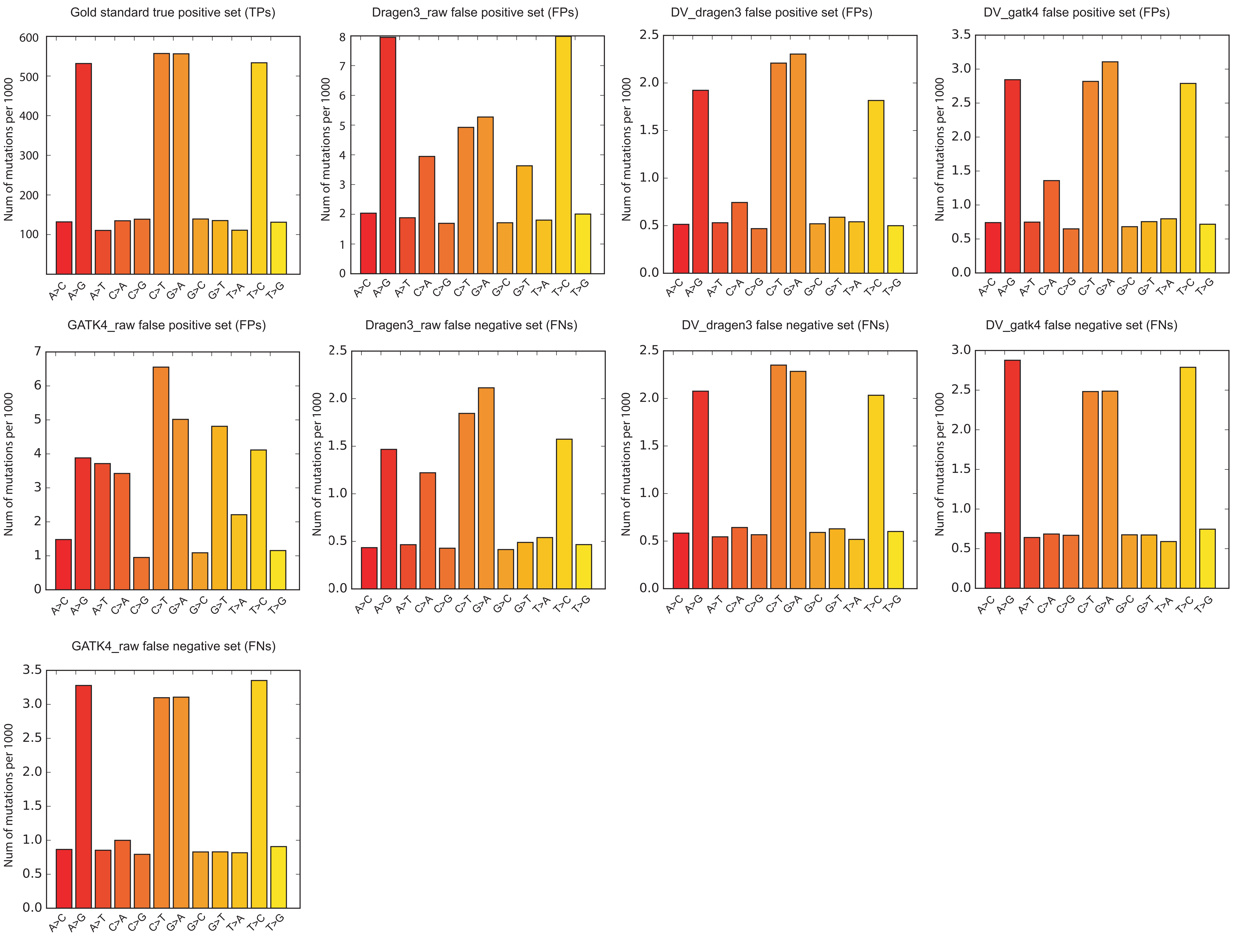

**Figure S4B.** The distribution of substitution signature of false positive and negative variants for NA12878_PrecisionFDA dataset

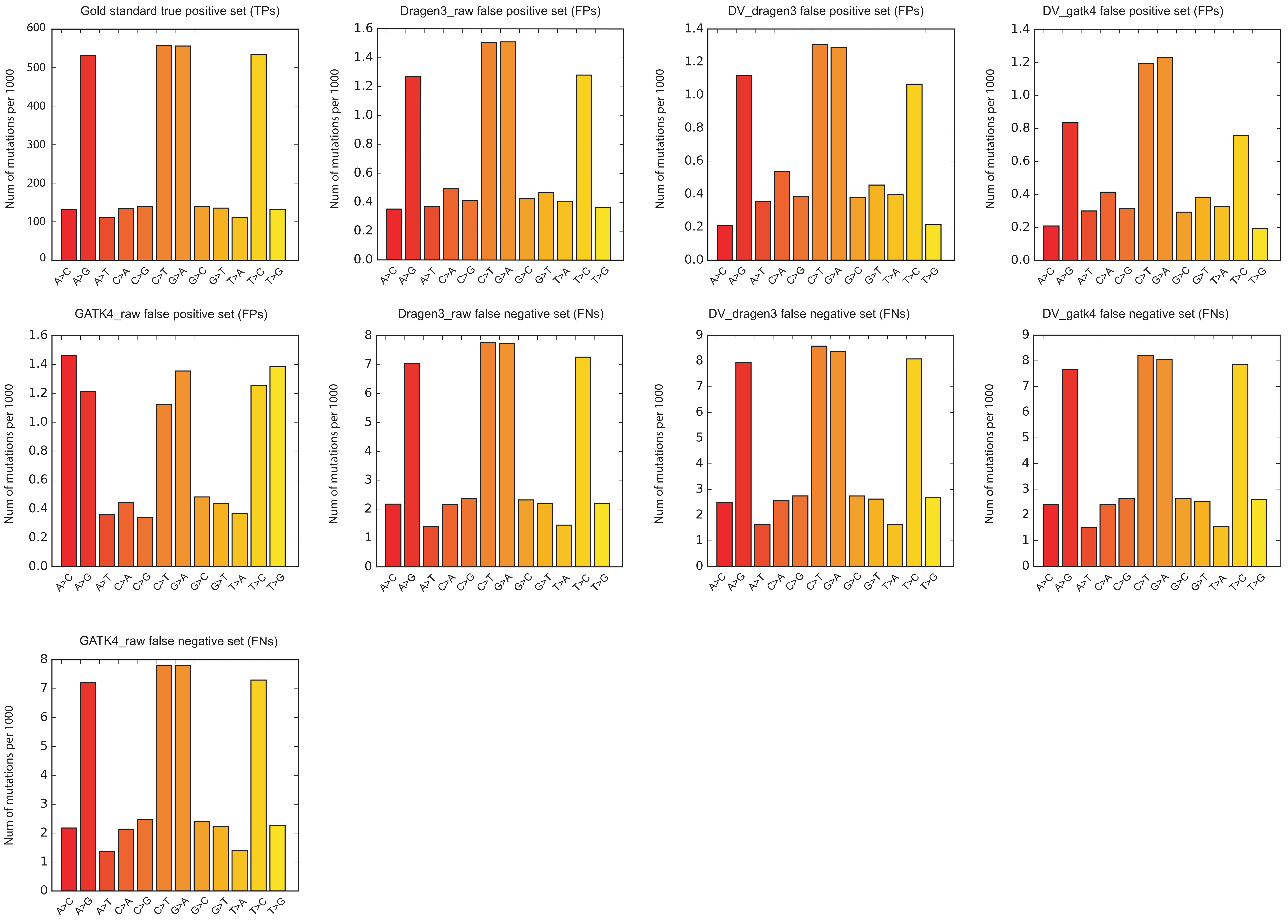

**Figure S4C.** The distribution of substitution signature of false positive and negative variants for the “simulated” dataset (random mutation profile)

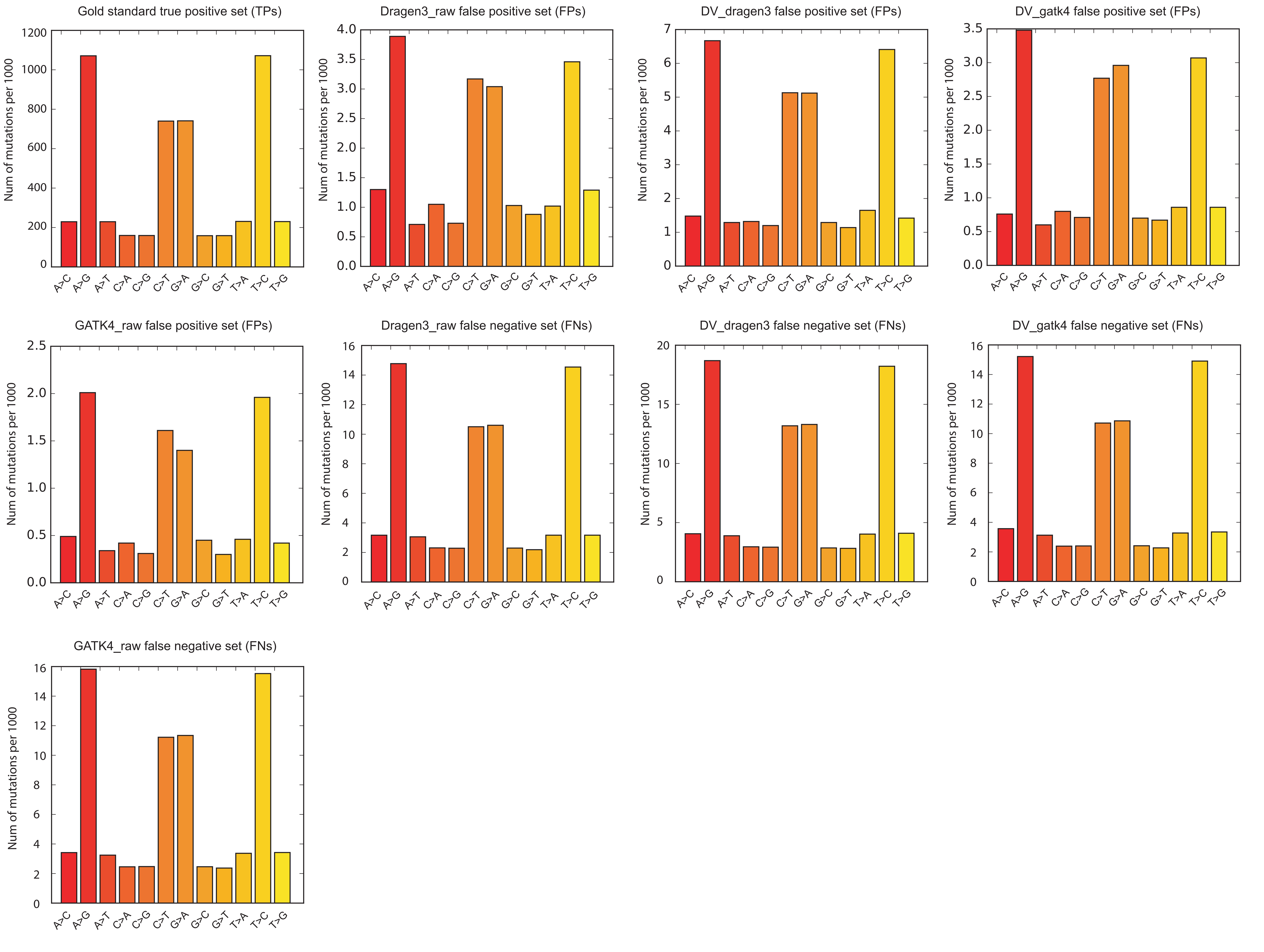

**Figure S4D.** The distribution of substitution signature of false positive and negative variants for the “simulated” dataset (userdefined mutation profile)

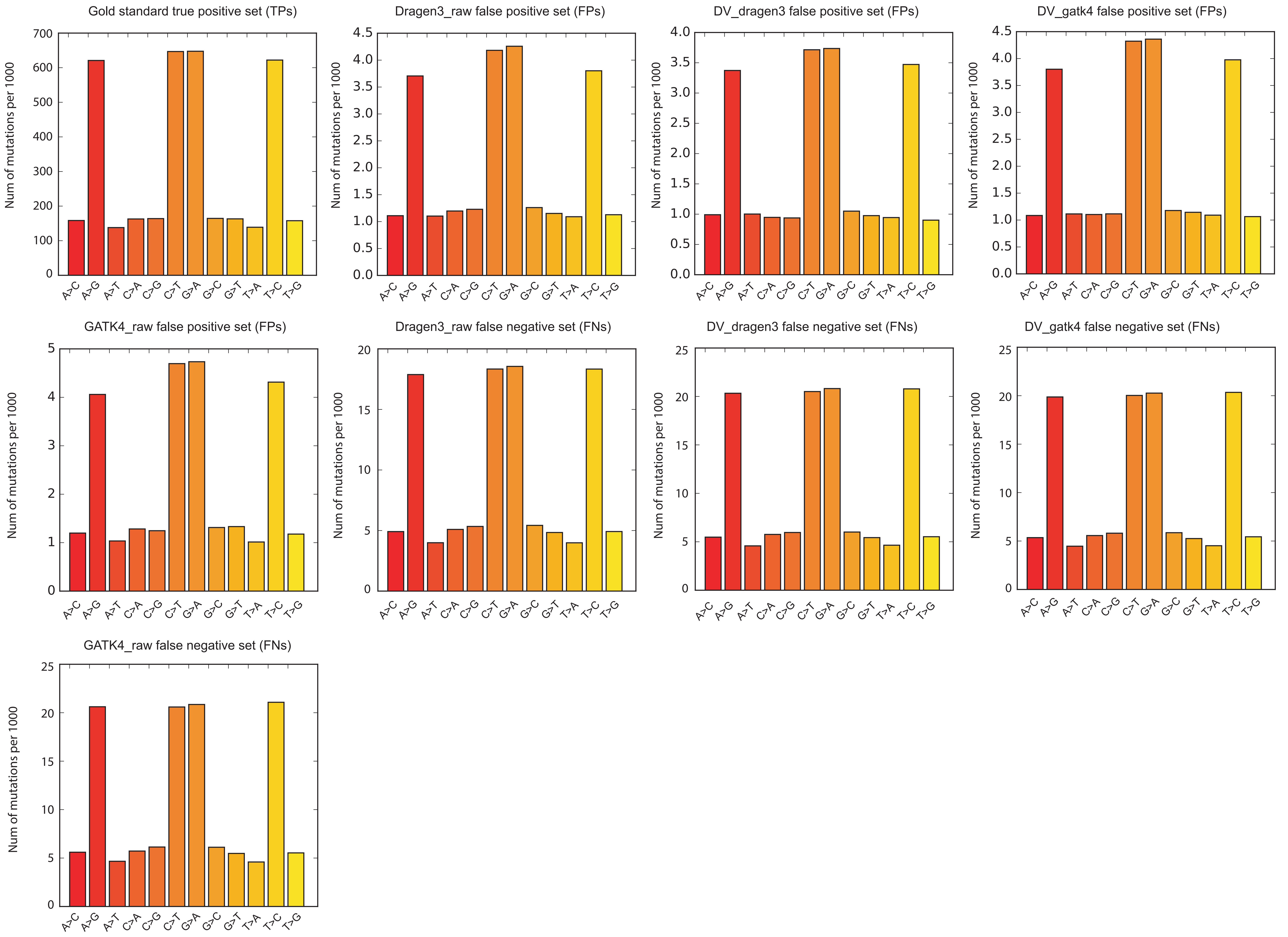

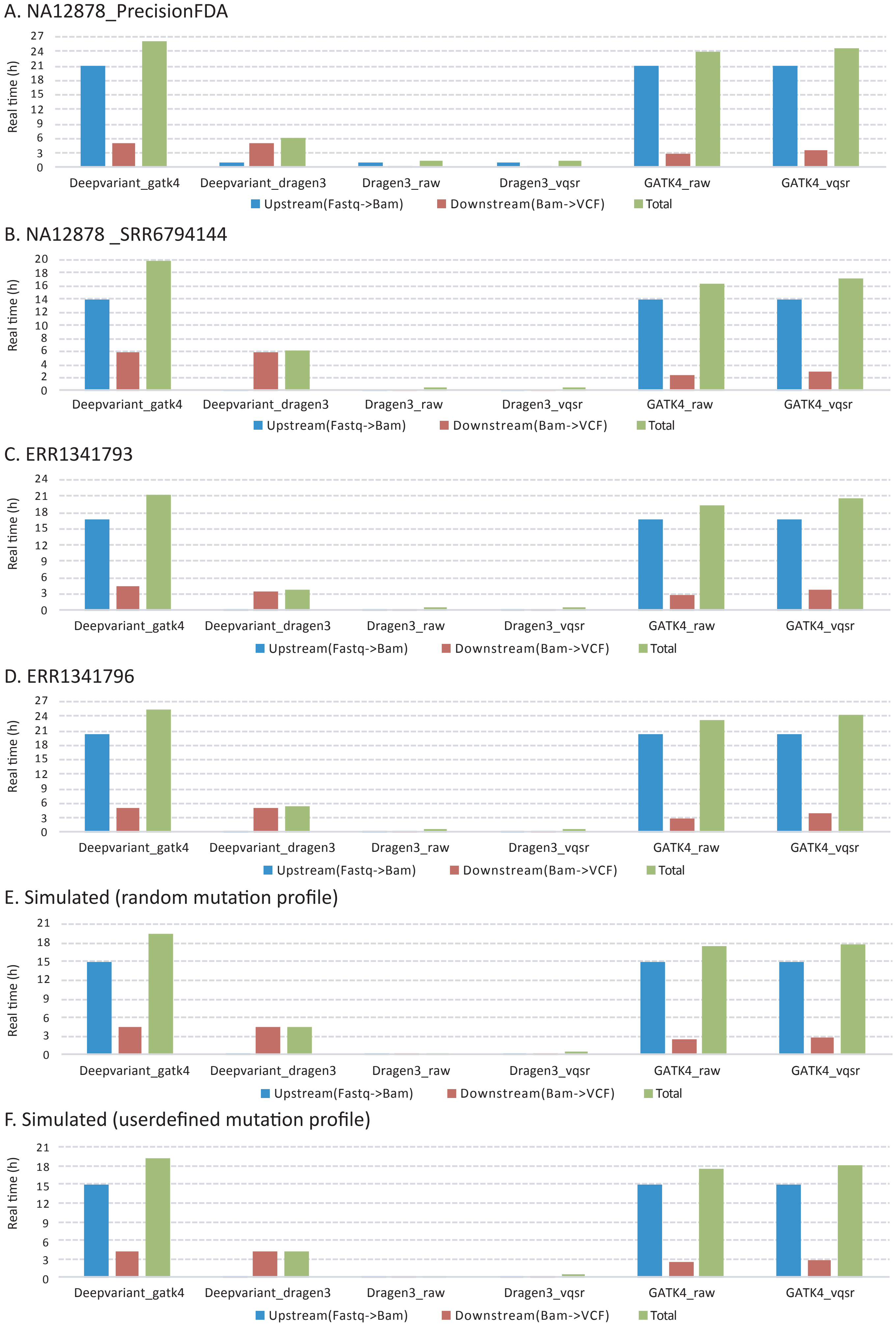


**Figure S5.** Variant calling runtime of six pipeline combinations (*DV_gatk4, DV_dragen3, Dragen3_raw, Dragen3_vqsr, GATK4_raw* and *GATK4_vqsr*) benchmarked on local virtual machine (VM). (A and B – NA12878_PrecisionFDA and NA12878_SRR6794144 datasets; C and D – "synthetic-diploid" ERR1341793 and ERR1341796 datasets; E and F - simulated data based on a random and a userdefined mutation profiles).
